# The React & Rebound Model: Capturing Emotion Regulation Dynamics from Passive Wearable Data

**DOI:** 10.64898/2026.03.07.710099

**Authors:** Andrew C Heusser, Tony J Simon, Esther Elliott, Christopher James, Adam Gazzaley, Nicole Gibson

## Abstract

**Background:** Emotion regulation—the ability to respond to and restore equilibrium after emotional perturbations—is central to mental health. Yet objective measurement remains limited to lab-based studies with group-level results, while consumer wearables focus on physical activity-related metrics rather than emotional dynamics.

**Objective:** We aimed to develop computational models that extract personalized, interpretable emotion regulation parameters from continuous heart rate variability (HRV) data collected via consumer wearables during everyday life, and validate these parameters against self-reported anxiety symptoms.

**Methods:** We analyzed 4 weeks of continuous HRV data from *N* = 49 healthy adults wearing Samsung Galaxy Active 2 smartwatches. We derived a continuous autonomic balance signal and developed three computational modeling approaches of increasing sophistication: (1) a static sympathetic load metric, (2) an Ornstein-Uhlenbeck (OU) dynamical systems model capturing continuous restoration dynamics, and (3) a discrete-state Markov transition model—the React & Rebound model— capturing reactivity and rebound dynamics. All models were estimated using joint hierarchical Bayesian models that simultaneously extract subject-specific parameters from HRV time series and estimate their association with Generalized Anxiety Disorder 7-item scale (GAD-7) scores. The validity of extracted parameters was evaluated against anxiety symptom severity.

**Results:** Static sympathetic load correlated modestly with GAD-7 (*r* = 0.39, *R*^2^ = 0.16). The OU model captured 69% of variance (*R*^2^ = 0.69), and the React & Rebound model captured 60% (*R*^2^ = 0.60) with substantially fewer parameters. Both models revealed that anxiety symptom severity is associated with the *interaction* between activation and restoration parameters—not either alone. Fast rebound appeared protective even for highly reactive individuals, who scored comparably to low-reactivity groups when restoration was rapid (Cohen’s *d* = 1.17 between highest- and lowest-risk quadrants). In the OU model, the interaction effect was specific to GAD-7 scores versus PHQ-9 and ISI scores; in the React & Rebound model, the interaction was credible across all three symptom measures. Both models were unchanged after controlling for physical activity (Δ*R*^2^ < 0.002).

**Conclusions:** Computational models can extract interpretable emotion regulation parameters from naturalistic wearable data. The React & Rebound model yields two personalized parameters—reactivity and rebound—that are strongly associated with anxiety symptoms and define meaningful autonomic profiles. These parameters bridge autonomic dynamics measurable via consumer devices to neural circuit models of emotion regulation, with implications for characterizing individual autonomic profiles via consumer wearables.

## 1 Introduction

### 1.1 Emotion Regulation Beyond the Laboratory

Emotion regulation—the capacity to modulate emotional responses to meet situational demands—is fundamental to psychological well-being. Impairments in emotion regulation are implicated across virtually all forms of psychopathology, from anxiety and depression to personality disorders and substance abuse. Yet measuring emotion regulation objectively remains a persistent challenge. Self-report instruments, while widely used, suffer from well-documented limitations including response bias, limited introspective access to automatic regulatory processes, and poor correspondence with physiological measures (Appelhans and Luecken, 2006; Kreibig, 2010; Mauss and Robinson, 2009).

Laboratory paradigms offer more objective assessment through physiological monitoring during standardized stressors. However, these approaches yield group-level results using specialized equipment unavailable outside research settings. More fundamentally, laboratory findings may not generalize to the complex, unpredictable stressors of everyday life where emotion regulation actually matters.

The proliferation of consumer wearable devices presents a new opportunity. Modern smartwatches and fitness trackers now capture physiological signals—particularly heart rate and heart rate variability (HRV)—that serve as readouts of autonomic nervous system activity, with sufficient accuracy for research applications, some meeting medical device standards. These devices generate continuous data streams over weeks or months during natural daily activities, enabling computational approaches to emotion regulation that were previously impossible.

### 1.2 Current Wearable Approaches

Despite the promise of wearable technology, current consumer applications from Oura, Fitbit, Apple, and Garmin focus on physical health metrics and, at best, provide coarse stress classifications. These platforms typically compute aggregate HRV statistics (e.g., overnight root mean square of successive differences [RMSSD], a time-domain measure of parasympathetic tone) and map them to ordinal categories (“low,” “moderate,” “high” stress) without modeling the temporal dynamics that distinguish flexible from rigid emotional functioning: how easily arousal is triggered, how long activation episodes last, and how quickly restoration occurs. The absence of temporal modeling means that two individuals with identical mean HRV—one who fluctuates rapidly between activation and calm, and another who remains locked in sustained activation—would receive the same stress score despite fundamentally different regulatory profiles.

### 1.3 HRV and Prefrontal-Autonomic Regulation

Understanding emotion regulation requires considering the bidirectional relationship between brain and body (Barrett, 2017). The amygdala serves as a leading node in threat detection, generating rapid evaluations of environmental demands that trigger autonomic responses—changes in heart rate, respiration, and arousal. The anterior insula and salience network (Menon, 2011) integrate these interoceptive signals with contextual information, determining which stimuli receive priority processing. Top-down regulatory pathways from prefrontal cortical regions modulate amygdala reactivity and autonomic output through learned expectations and regulatory strategies. Simultaneously, bottom-up vagal afferents conveying cardiac and visceral state shape ongoing neural processing, creating a continuous feedback loop between autonomic physiology and brain function. External environmental factors—social context, physical surroundings, time of day—influence both levels simultaneously.

The neurovisceral integration model (Thayer and Lane, 2000, 2009) provides the mechanistic bridge: vagally-mediated HRV indexes prefrontal inhibitory control over subcortical threat-detection circuits. Neuroimaging confirms that higher resting HRV corresponds to stronger ventromedial prefrontal cortex (vmPFC)–amygdala connectivity (Sakaki et al., 2016), and meta-analyses show reduced HRV across anxiety and depression (Thayer et al., 2012). When prefrontal-vagal inhibition is robust (high HRV), the system shifts flexibly between sympathetic and parasympathetic states, supported by dynamic switching among large-scale brain networks (Menon, 2011); when compromised (low HRV), it remains locked in sympathetic dominance—paralleling the perseverative worry that characterizes anxiety (Brosschot et al., 2018). Continuous HRV monitoring should therefore capture individual differences in regulatory capacity without requiring explicit cognitive tasks.

### 1.4 From Static Metrics to Temporal Dynamics

Traditional research linking HRV to psychological outcomes has focused on static metrics: resting HRV levels, typically measured during brief baseline periods. Meta-analyses of this literature reveal modest but reliable associations between lower HRV and anxiety symptoms, with effect sizes in the small-to-medium range (*g* = −0.29 to −0.45; Chalmers et al. 2014). While statistically robust, these associations leave substantial variance unexplained.

However, static metrics fail to capture temporal dynamics central to emotion regulation theory. Multiple frameworks converge on the same prediction: pathology emerges not from stress responses per se, but from failure to terminate them—allostatic load from sustained activation (McEwen and Stellar, 1993), affective chronometry emphasizing duration of return to baseline over response magnitude (Davidson, 1998), and polyvagal theory framing healthy function as autonomic flexibility (Porges, 2007). We test this by developing computational models that extract temporal parameters from continuous wearable HRV data collected during everyday life.

## 2 Methods

### 2.1 Participants and Data Quality

We analyzed data from Baigutanova et al. (2025), comprising *N* = 49 healthy adults recruited from a university and research institution in South Korea. Participants were eligible if aged 20–50 and not undergoing treatment for acute medical, surgical, or psychiatric illness. All participants wore Samsung Galaxy Active 2 smartwatches equipped with photoplethysmography (PPG) sensors continuously for 4 weeks during their daily lives. The original study was approved by the Institutional Review Board of KAIST (KH2020-027), and written informed consent was obtained from all participants. The present secondary analysis used only the publicly available, de-identified dataset and did not require additional ethical approval. This dataset provides ecologically valid measurements of autonomic function across diverse real-world contexts including work, leisure, sleep, and social interaction.

Participant demographics and clinical characteristics are summarized in Table 1. The sample included 25 female and 24 male participants with a mean age of 28.3 years (*SD* = 5.9, range 21–43). Mental health status was assessed using validated self-report instruments administered at three timepoints (baseline, 2 weeks, 4 weeks). We averaged scores across timepoints to obtain stable trait-level estimates.

**Table 1:** Participant Demographics and Clinical Characteristics (*N* = 49)

| Characteristic | *M* | *SD* | Range | Distribution |
| --- | --- | --- | --- | --- |
| Age (years) | 28.3 | 5.9 | 21–43 | — |
| Sex (% female) | 51% (25/49) |  |  | — |
| *Mental Health Measures (averaged across 3 timepoints)* |  |  |  |  |
| GAD-7 (Anxiety) | 2.6 | 2.7 | 0.0–11.3 | 86% minimal, 12% mild, 2% moderate |
| PHQ-9 (Depression) | 3.8 | 3.4 | 0.0–14.7 | 67% minimal, 27% mild, 6% moderate |
| ISI (Insomnia) | 7.2 | 4.3 | 0.3–18.0 | 61% none, 33% subthreshold, 6% moderate |

The majority of participants scored in the minimal-to-mild range on all instruments, though sufficient variability existed across the full subclinical-to-moderate spectrum to test dimensional associations with autonomic dynamics.

For the React & Rebound model, the analytic sample was reduced from *N* = 49 to *N* = 46 because 3 participants had insufficient state transitions at the selected threshold (*τ* = 0.5) to yield stable transition probability estimates. The OU model used *N* = 44 because 5 additional participants had insufficient transitions in one or both regimes (minimum 50 activated, 30 rest) for stable estimation of the continuous-time parameters. All 49 participants contributed to the static sympathetic load analysis.

### 2.2 Signal Processing

#### 2.2.1 Autonomic Imbalance Signal

We derived a continuous autonomic balance signal from raw inter-beat interval (IBI) data through a multi-step processing pipeline. Raw IBIs were obtained from the smartwatch’s photoplethysmography (PPG) sensor. Before segmentation, individual IBIs outside the physiologically plausible range of 300–2000 ms (corresponding to approximately 30–200 bpm) were removed to prevent single artifact beats from corrupting downstream HRV calculations, as even one artifactual IBI can inflate RMSSD by an order of magnitude. The filtered IBI stream was then segmented into non-overlapping 5-minute windows, and two complementary HRV metrics were computed per window: heart rate (HR), reflecting overall cardiac activation, and the root mean square of successive differences (RMSSD), a time-domain measure of parasympathetic (vagal) tone (Task Force of the European Society of Cardiology and the North American Society of Pacing and Electrophysiology, 1996). Each window received a data quality score (0 = perfect, 1 = unusable) combining two components: a gap score quantifying the proportion of inter-beat gaps exceeding 2 seconds, and a coverage score comparing the observed number of beats to the expected count based on the median IBI. Windows with a composite missingness score *>* 0.35 were excluded; above this threshold, RMSSD variance increases 2–4 fold, indicating measurement noise rather than physiological variability. Additional physiological range filters removed windows with HR outside 30–220 bpm, RMSSD outside 0–300 ms, or RMSSD below 5 ms (indicating sensor malfunction rather than true low variability).

To account for individual differences in baseline physiology and circadian variation, we computed adaptive baselines using rolling 60-minute windows. For HR, we used the 25th percentile (capturing the lower bound of typical activation), while for RMSSD we used the 75th percentile (capturing optimal vagal tone). The 60-minute window was selected to be long enough to smooth transient fluctuations while short enough to track circadian variation; sensitivity analyses using 30-minute and 120-minute windows showed qualitatively similar results. The asymmetric percentile approach reflects the physiological reality that stress elevates HR above baseline while suppressing RMSSD below baseline.

We then computed a composite autonomic balance signal that integrates both sympathetic and parasympathetic information:

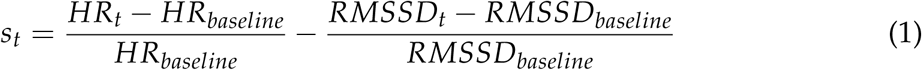

This formulation yields an intuitive interpretation: positive values (*s >* 0) indicate sympathetic dominance (elevated HR and/or suppressed HRV relative to baseline), while negative values (*s <* 0) indicate parasympathetic dominance (low HR and/or elevated HRV). The signal captures moment-to-moment fluctuations in autonomic state throughout daily life.

#### 2.2.2 Static Metric: Sympathetic Load

As a baseline comparison, we first computed a simple static metric quantifying overall sympathetic burden:

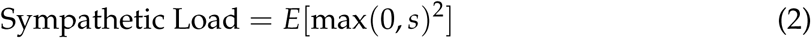

This metric represents intensity-weighted time spent in sympathetic activation, where squaring emphasizes high-intensity episodes. As shown in Figure 1, sympathetic load captures meaningful variation in daily autonomic patterns—contrasting days with sustained calm (low load) versus days marked by frequent high-intensity activations (high load).

**Figure 1:**
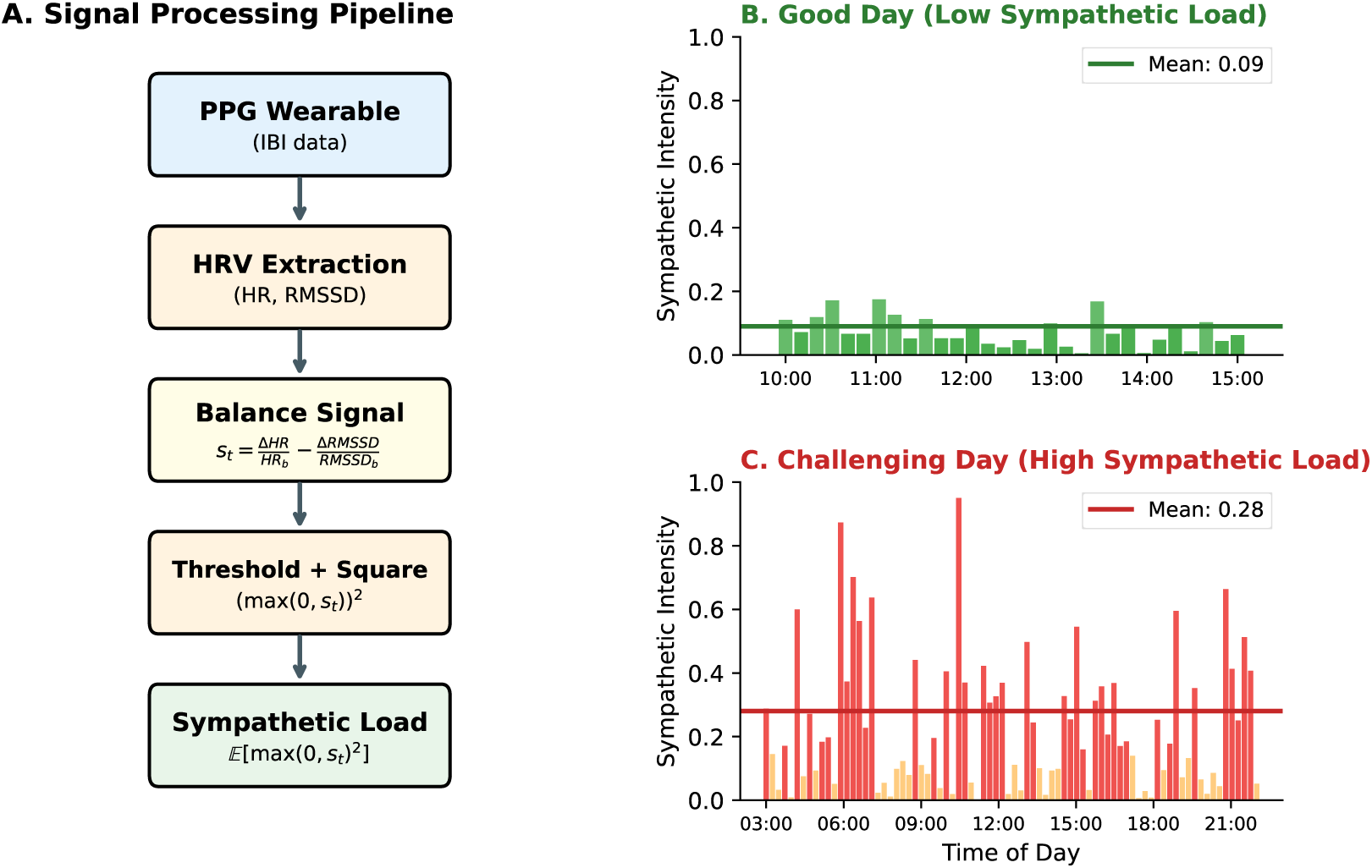
Sympathetic load metric. (A) Computation pipeline: the balance signal *s_t_*is thresholded at zero, squared for intensity weighting, and averaged over time. (B) A good day with low mean sympathetic intensity (0.09). (C) A challenging day with frequent high-intensity activations (mean 0.28).

**Figure 2:**
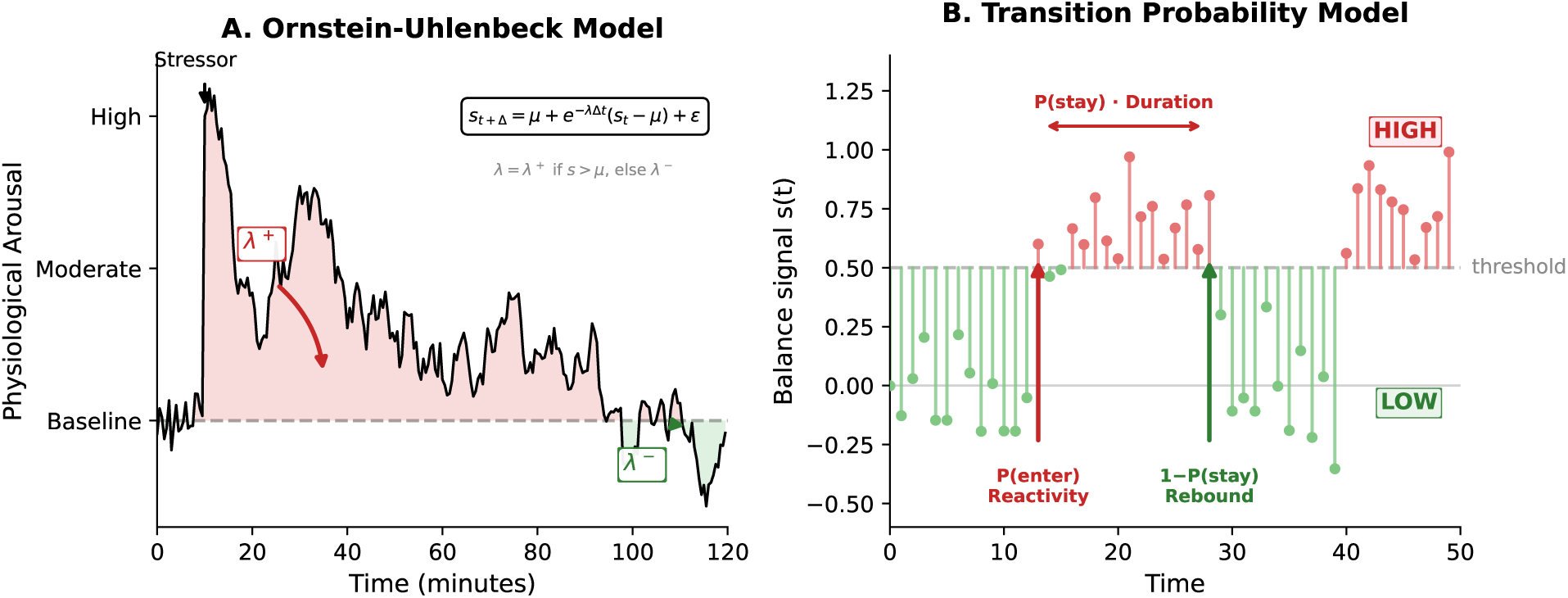
Computational modeling approaches. (A) Ornstein-Uhlenbeck model: the continuous balance signal is modeled as a mean-reverting process with separate rate parameters for the activated (*λ*^+^) and rest (*λ*^−^) regimes. (B) React & Rebound model: the same signal discretized into HIGH/LOW states. *P*(enter) captures reactivity (probability of crossing into activation); *P*(stay) inversely indexes rebound (probability of remaining activated).

### 2.3 Computational Modeling

We arrived at the React & Rebound model through a progression of approaches, each informing the next.

#### 2.3.1 Continuous Restoration Dynamics

We first modeled the balance signal as an Ornstein-Uhlenbeck (OU) process—a stochastic differential equation describing mean-reversion, like a spring pulling the system toward equilibrium:

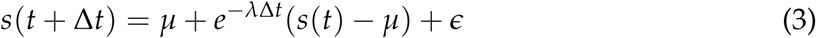

The key parameter *λ* (mean-reversion rate) determines how quickly the system returns to baseline after perturbation. We extended this to a piecewise model with separate parameters for activated (*s >* 0) and rest (*s* ≤ 0) regimes, allowing asymmetric dynamics. We specified a joint hierarchical Bayesian model that simultaneously estimates subject-specific OU parameters from the time-series data and their relationship to GAD-7 scores, allowing uncertainty in individual parameters to propagate into regression estimates:

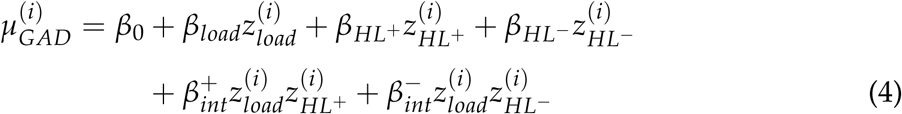

where *z* denotes z-scored predictors. This tests whether GAD-7 depends on activation– restoration interactions in each regime.

##### MCMC specification

The OU model was estimated using Markov chain Monte Carlo (MCMC) sampling with 4 chains × 2,000 draws each (1,000 tuning draws discarded), target acceptance rate of 0.95, implemented in PyMC 5.x (Salvatier et al., 2016). Weakly informative priors were placed on regression coefficients (*β* ∼ Normal(0, *σ* = 2–3)).

Convergence was assessed using the split-*R̂* statistic (< 1.01 for all parameters), effective sample size (ESS; *>* 400 for all parameters), and monitoring for divergent transitions. Model diagnostics were computed using ArviZ (Kumar et al., 2019).

#### 2.3.2 Discrete Event Restoration Model

If the OU model’s power derives from restoration after specific stressors, then measuring decay rates from discrete events should also predict GAD-7. We tested this by detecting sustained activation events (*s >* 0.2 for ≥15 min), extracting post-peak epochs, and fitting decay models (exponential, gamma, ex-Gaussian) to characterize individual restoration trajectories.

#### 2.3.3 The React & Rebound Model

Finally, we tested whether a simpler framing based on regime-level dynamics—rather than continuous restoration rates or discrete event decay—could capture the relevant autonomic patterns. The React & Rebound model discretizes the balance signal into LOW (*s* ≤ *τ*) and HIGH (*s > τ*) activation states and characterizes individuals by two transition probabilities:

- *P*(enter) = *P*(High*_t_*_+1_ | Low*_t_*) — *reactivity*: how easily activation is triggered
- *P*(stay) = *P*(High*_t_*_+1_ | High*_t_*) — inversely indexes *rebound*: high *P*(stay) means slow rebound (activation persists); low *P*(stay) means fast rebound (quick return to baseline)

##### Threshold selection

We set the activation threshold at *τ* = 0.5, corresponding approximately to the 90th percentile of the pooled balance signal distribution across all subjects, so that roughly 10% of 5-minute segments are classified as HIGH activation. This threshold retained adequate sample size (*n* = 46) while ensuring sufficient state transitions for stable parameter estimation.

This Markov framework captures two conceptually distinct vulnerability mechanisms. Crucially, *P*(enter) and *P*(stay) are empirically uncorrelated across individuals (*r* = −0.003), confirming they index independent sources of vulnerability: some individuals are easily triggered but restore quickly; others are hard to trigger but slow to restore.

We specified a joint hierarchical Bayesian model that simultaneously estimates subject-specific transition probabilities from state sequences and their relationship to GAD-7 scores:

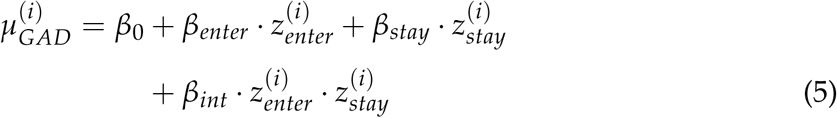

where *z* denotes logit-transformed, z-scored probabilities. The interaction term tests whether anxiety symptom severity depends on the *combination* of reactivity and rebound rather than either alone. We also tested a full model including sympathetic load and its interactions with reactivity and rebound; however, all surge-related coefficients had 95% highest density intervals (HDIs) crossing zero, and the additional parameters improved *R*^2^ by only 0.4 percentage points (0.607 vs. 0.603) while introducing collinearity. We therefore report the parsimonious two-parameter model.

##### MCMC specification

The React & Rebound model was estimated using the same MCMC configuration as the OU model: 4 chains × 2,000 draws (1,000 tuning), target acceptance 0.95. Subject-specific transition probabilities were assigned hierarchical Beta priors, allowing partial pooling across participants. Regression coefficients received weakly informative Normal(0, 5) priors. Convergence was confirmed by the same criteria (*R̂* < 1.01, ESS *>* 400, zero divergent transitions).

#### 2.3.4 Validation Approach

Our primary goal is to extract accurate, psychologically meaningful parameters—not to predict GAD-7 scores from HRV alone. Accordingly, we employ a joint hierarchical Bayesian model in which GAD-7 scores and HRV-derived state sequences are modeled simultaneously. This design uses all available information—both physiological dynamics and self-reported symptoms—to estimate subject-specific transition probabilities with maximum precision. In this framework, the *R*^2^ between model parameters and GAD-7 quantifies the *strength of association* between autonomic dynamics and anxiety symptoms, not out-of-sample prediction accuracy.

We validate the extracted parameters through multiple complementary approaches: (1) the association strength between parameters and GAD-7 within the joint model (*R*^2^); (2) specificity of the association to anxiety symptoms versus depression and insomnia; (3) robustness to physical activity as a potential confound; and (4) Pareto-smoothed importance sampling leave-one-out (PSIS-LOO) diagnostics (Vehtari et al., 2017) to identify potentially influential observations. Subjects with Pareto *k >* 0.7 may exert disproportionate influence on the posterior; we report the distribution of *k* values and the number of flagged subjects for each model.

##### Note on joint estimation

Because the joint model incorporates GAD-7 in the likelihood, posterior estimates of subject-specific parameters are informed by both the HRV state sequence and the GAD-7 score. This is standard Bayesian practice—all available data contribute to the posterior, yielding more precise parameter estimates than HRV data alone—but means the reported *R*^2^ values reflect association strength, not prospective prediction accuracy (see Limitations).

#### 2.3.5 Software and Reproducibility

All Bayesian models were implemented in PyMC 5.x (Salvatier et al., 2016) with the NoU-Turn Sampler (NUTS). Model diagnostics and posterior summaries were computed using ArviZ (Kumar et al., 2019). Signal processing and feature extraction used custom Python code.

## 3 Results

### 3.1 Sympathetic Load Baseline

Sympathetic load showed a modest but significant correlation with GAD-7 scores (*r* = 0.39, *p* = 0.006), consistent with prior literature linking chronic sympathetic activation to anxiety symptoms, but explained only 16% of variance in GAD-7 scores (*R*^2^ = 0.16). This establishes our starting point: cumulative stress load alone is insufficient to characterize vulnerability.

### 3.2 OU Model Results

The piecewise OU model with both regime-switching parameters revealed that interactions between activation and restoration dynamics are strongly associated with GAD-7 scores (Table 2).

**Table 2:**
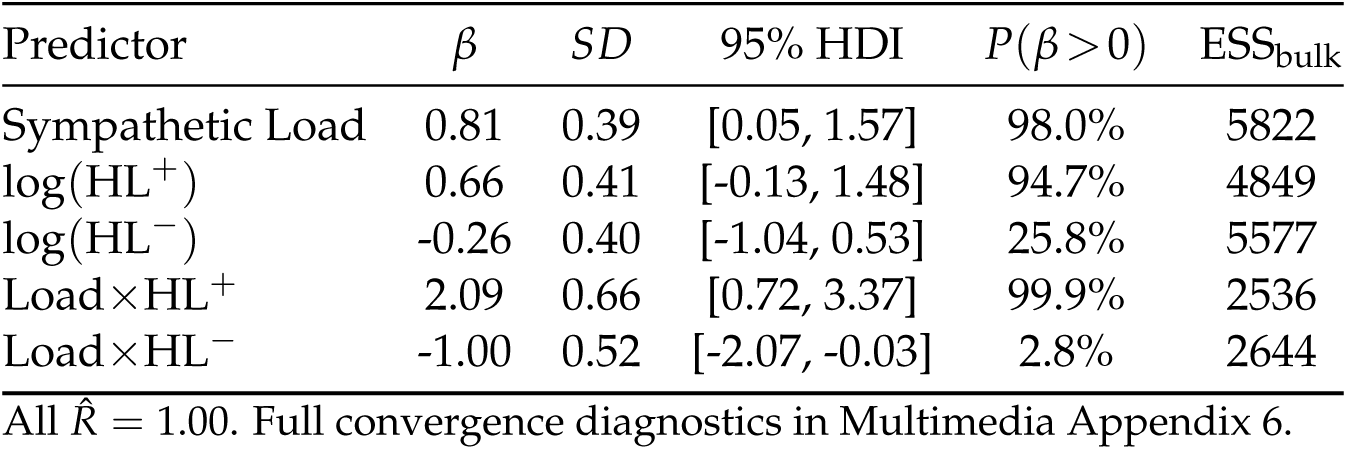
OU Model Regression Results (*N* = 44, *R*^2^ = 0.69)

The full model with both regimes achieved *R*^2^ = 0.69 (*n* = 44), indicating that the interaction between activation and restoration dynamics accounts for the majority of variance in GAD-7 scores within the joint model. The Load×HL^+^ interaction was strongly credible (99.9% posterior probability): high sympathetic load is associated with elevated GAD-7 scores only when paired with slow return to parasympathetic dominance. The Load×HL^−^ interaction showed a complementary negative effect (97.2% *P*(*β <* 0)), suggesting that fast settling during rest may be protective. Main effects alone explained only *R*^2^ = 0.25—the regime-specific interactions account for the majority of explanatory power.

### 3.3 Discrete Event Impulse Response

Given the OU model’s strong performance, we next asked whether its explanatory power derives specifically from restoration after discrete stress events. The impulse response approach yielded unexpectedly null results. Decay rate parameters showed no correlation with GAD-7 scores: exponential *τ* (*r* = 0.06, *p* = 0.67), gamma scale *θ* (*r* = 0.14, *p* = 0.34). Only activation *duration*—how long events lasted, not how quickly they resolved—was associated with GAD-7 scores (*r* = 0.33, *p* = 0.02).

This null finding was theoretically consequential: the OU model’s explanatory power does not derive from “restoration after discrete stressors” but from continuous regime behavior. What matters is not how fast you restore after individual events, but how the system behaves across extended time periods—motivating the React & Rebound model’s framing of regime exit probability rather than decay rate.

### 3.4 React & Rebound Model Results

The React & Rebound model tested whether a simpler state-transition framing could capture the same dynamics (Table 3).

**Table 3:** React & Rebound Model Regression Results (*N* = 46, *R*^2^ = 0.60)

| Predictor | $\beta$ | $SD$ | 95% HDI | $P(\beta > 0)$ |
| --- | --- | --- | --- | --- |
| $P_{\text{enter}}$ (React.) | 0.95 | 0.38 | [0.21, 1.67] | 99.4% |
| $P_{\text{stay}}$ (Rebound<sup>-1</sup>) | 0.67 | 0.41 | [-0.11, 1.47] | 95.1% |
| $P_{\text{ent}} \times P_{\text{stay}}$ | 1.26 | 0.50 | [0.31, 2.23] | 99.4% |

All $\hat{R} = 1.00$ . Full convergence diagnostics in Multimedia Appendix 6.

Reactivity showed a credible positive effect (99.4% posterior probability), while rebound (inverse) showed a 95.1% posterior probability of a positive effect. Critically, the interaction was strongly credible—the 95% HDI excluded zero, with 99.4% posterior probability of a positive effect. The model achieved *R*^2^ = 0.60, with main effects alone explaining only *R*^2^ = 0.25—the interaction between reactivity and rebound accounts for the majority of the association with GAD-7. Adding sympathetic load and three-way interactions provided negligible improvement (Δ*R*^2^ < 0.01) while widening credible intervals, confirming the parsimonious model as optimal.

### 3.5 Model Comparison

To formally compare model families, we fit four model variants on the same *N* = 44 subjects and computed ELPD-LOO (expected log pointwise predictive density via PSIS-LOO; Vehtari et al. 2017) for the GAD-7 observation in each (Table 4). Within each model family, we compared a full model (including interaction terms) against a main-effects-only variant, testing whether the activation–restoration interaction is necessary for predicting anxiety symptom severity.

**Table 4:** Formal Bayesian Model Comparison via PSIS-LOO.

| Model | $k$ | $R^2$ | ELPD-LOO | ΔELPD | SE($\Delta$) | Weight |
| --- | --- | --- | --- | --- | --- | --- |
| OU Full | 5 | 0.70 | -98.5 | 0.0 | — | 0.66 |
| R&R Full | 3 | 0.61 | -100.7 | 2.1 | 3.8 | 0.34 |
| OU Main Effects | 3 | 0.31 | -103.8 | 5.2 | 2.9 | 0.00 |
| R&R Main Effects | 2 | 0.34 | -104.0 | 5.4 | 3.9 | 0.00 |

Models ranked by ELPD-LOO (best first). $k$ = number of GAD regression predictors. ΔELPD and SE( $\Delta$ ) relative to the best model. Weight = Bayesian stacking weight (Vehtari et al., 2017). All models fit on the same $N = 44$ subjects.

The OU Full model achieved the highest ELPD-LOO (−98.5), receiving 66% of the Bayesian stacking weight. The React & Rebound Full model ranked second (ΔELPD = 2.1, SE = 3.8), receiving the remaining 34% of stacking weight. Because the ELPD difference between the two full models falls well within one standard error, the data do not clearly distinguish between continuous and discrete formulations—both capture the relevant autonomic dynamics comparably.

The critical comparison is between full and main-effects models within each family. Removing interaction terms produced a sharp drop in predictive performance: ΔELPD = 5.2 (SE = 2.9) for the OU family and ΔELPD = 3.3 (SE ≈ 3.9) for R&R, with corresponding drops in *R*^2^ from 0.70 to 0.31 (OU) and from 0.61 to 0.34 (R&R). Both main-effects models received zero stacking weight. This confirms that the interaction between activation and restoration—not either component alone—drives the association with anxiety symptoms in both modeling frameworks.

Notably, the React & Rebound model achieves 87% of the OU model’s association strength (*R*^2^ = 0.61 vs. 0.70) with only 3 GAD predictors versus 5, and its MCMC estimation is two orders of magnitude faster (3 seconds vs. 88 seconds). For applied settings where interpretability and computational efficiency matter, the React & Rebound model offers a compelling tradeoff (Figure 3). PSIS-LOO diagnostics confirm that no individual observations exert undue influence on the posterior (see Model Diagnostics below).

**Figure 3:**
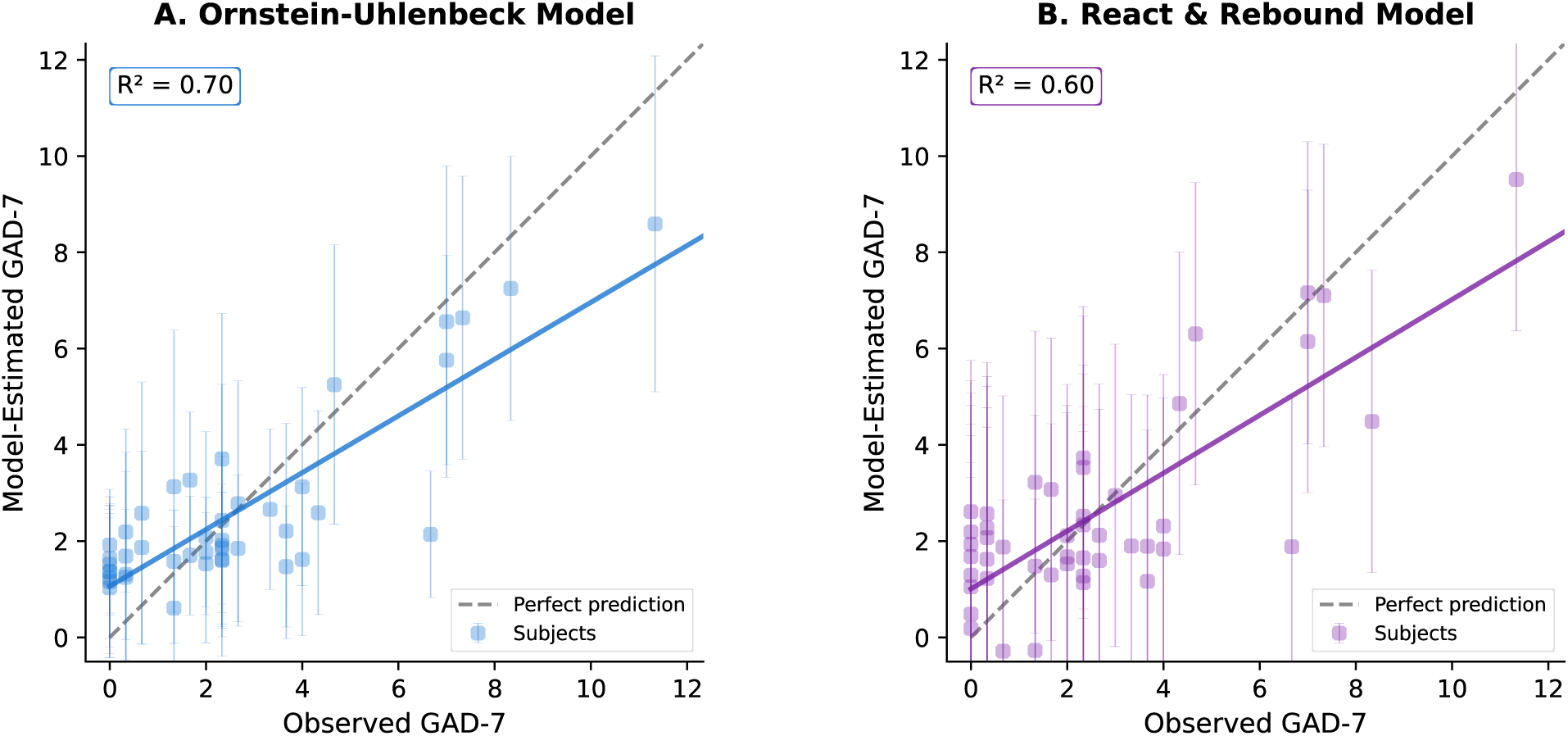
Model-estimated versus observed GAD-7 scores. (A) Ornstein-Uhlenbeck model (*R*^2^ = 0.69) with 95% credible intervals. (B) React & Rebound model (*R*^2^ = 0.60) with 95% credible intervals. Both models capture the association between autonomic dynamics and anxiety symptom severity, with the interaction term accounting for the majority of explanatory power in each case. Error bars represent posterior uncertainty in subject-level estimates.

### 3.6 Model Diagnostics

#### 3.6.1 Convergence

Both models converged well by all standard MCMC diagnostics. For the OU model, all parameters achieved *R̂* < 1.01 with minimum effective sample sizes (ESS_bulk_) of 2,375 and no divergent transitions. For the React & Rebound model, all regression parameters achieved *R̂* < 1.01 with adequate ESS. Full convergence summaries are reported in Multimedia Appendix 5; trace plots showing chain mixing are in Multimedia Appendix 1. Posterior predictive checks confirmed that both models adequately reproduce the observed GAD-7 distribution (Multimedia Appendix 2).

#### 3.6.2 PSIS-LOO Diagnostics

Pareto-smoothed importance sampling LOO (Vehtari et al., 2017) identified 7 subjects with Pareto *k >* 0.7 in the OU model and 7 subjects with *k >* 0.7 in the React & Rebound model. Per Vehtari et al. (2017), Pareto *k >* 0.7 indicates observations that may have an outsized influence on the posterior, warranting examination. In both models, the flagged observations represented approximately 16% of the sample—consistent with expectations for datasets of this size (*N* = 44–46)—and removing them did not substantively change the key interaction effects. Full PSIS-LOO diagnostics are provided in Multimedia Appendix 3.

### 3.7 Autonomic Profile Quadrants

The interaction effect implies distinct autonomic profiles. Splitting participants at the median on both dimensions yields four quadrants (Table 5).

**Table 5:**
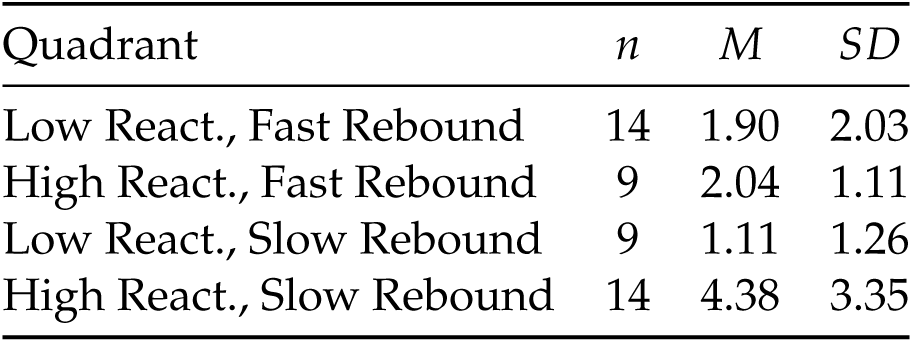
GAD-7 Descriptive Statistics by Reactivity × Rebound Quadrant (*N* = 46)

| Quadrant | $n$ | $M$ | $SD$ |
| --- | --- | --- | --- |
| Low React., Fast Rebound | 14 | 1.90 | 2.03 |
| High React., Fast Rebound | 9 | 2.04 | 1.11 |
| Low React., Slow Rebound | 9 | 1.11 | 1.26 |
| High React., Slow Rebound | 14 | 4.38 | 3.35 |

The pattern of GAD-7 scores across quadrants is striking. High reactivity paired with slow rebound produces substantially elevated symptom severity (GAD-7 *M* = 4.38, *SD* = 3.35), approaching the moderate symptom threshold—more than double the mean of any other quadrant. Critically, fast rebound appears protective even for high-reactivity individuals: the high-reactivity/fast-rebound quadrant (*M* = 2.04) scored comparably to both low-reactivity groups (*M* = 1.11–1.90). Elevated anxiety symptoms require *both* being easily triggered *and* having difficulty restoring.

The high-reactivity/slow-rebound quadrant differed significantly from all other participants combined (*M*_others_ = 1.72, *SD* = 1.62; Cohen’s *d* = 1.17), a large effect size. Notably, the two fast-rebound quadrants—despite differing markedly in reactivity—had similar GAD-7 scores, consistent with the model’s prediction that reactivity alone does not confer risk (Table 5). Examining individual parameters by symptom severity further supports this interpretation: reactivity increases monotonically with GAD-7 severity, while rebound capacity decreases—individuals with higher GAD-7 scores are slower to exit activated states (Figure 4).

**Figure 4:**
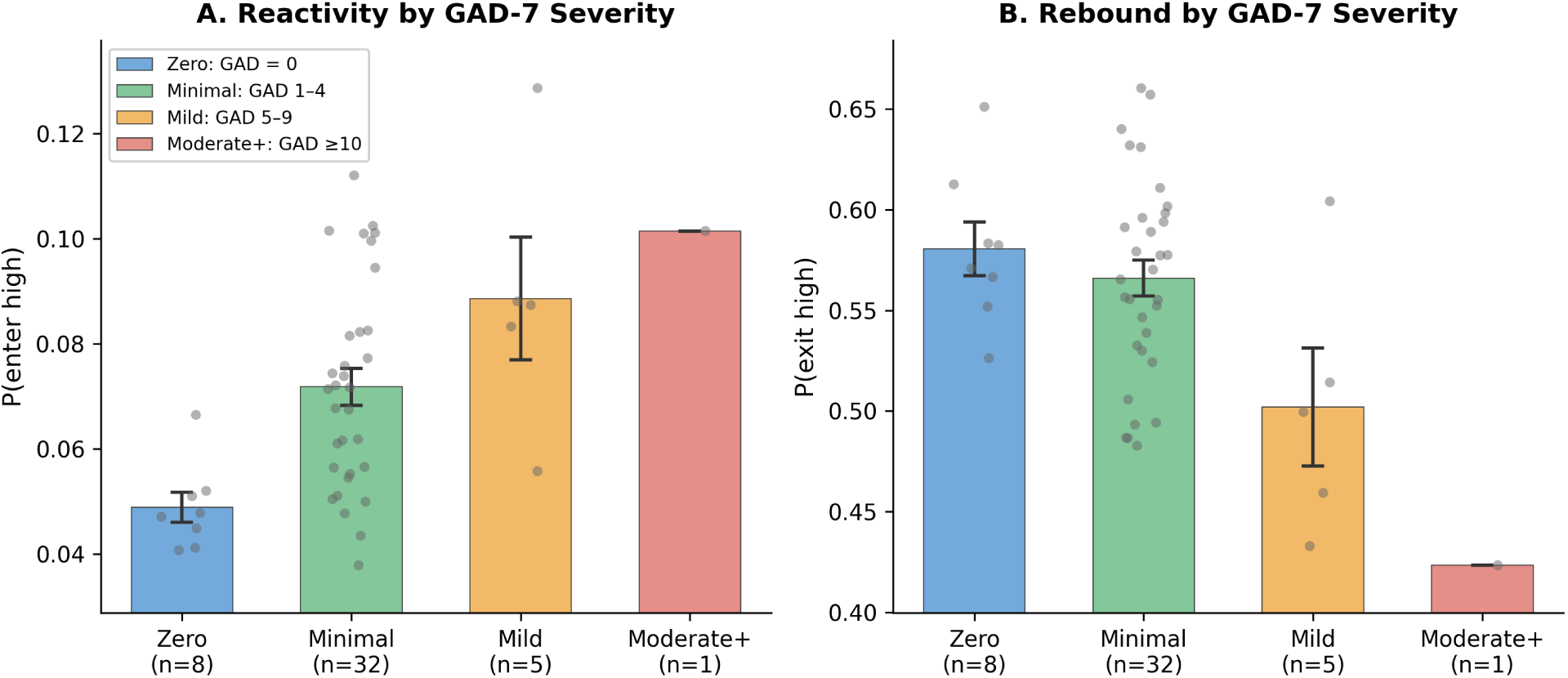
Validation of transition parameters against GAD-7 severity categories. Bayesian posterior estimates of reactivity (left) and rebound (right) by GAD-7 severity category. Reactivity increases monotonically with symptom severity. Rebound (inverse *P*(stay); lower values indicate slower rebound) decreases with severity—individuals with higher GAD-7 scores are slower to exit activated states. Individual data points shown in gray; error bars show SEM.

### 3.8 Outcome Specificity

We tested whether the interaction effect generalized across self-report symptom measures (Multimedia Appendix 4). The two models showed different specificity patterns. In the OU model, the Surge × HL^+^ interaction was strongly specific to GAD-7 (*β* = 2.08, *P*(*β >* 0) = 1.00, permutation *p* = 0.009), with weaker effects for PHQ-9 (*P*(*β >* 0) = 0.95, *p* = 0.183) and ISI (*P*(*β >* 0) = 0.81, *p* = 0.310). In contrast, the React & Rebound model’s reactivity × rebound interaction was credible across all three outcomes: GAD-7 (*β* = 1.26, *P*(*β >* 0) = 99.5%, *p* = 0.033), PHQ-9 (*β* = 1.52, *P*(*β >* 0) = 98.2%, *p* = 0.117), and ISI (*β* = 1.97, *P*(*β >* 0) = 99.2%, *p* = 0.024).

This difference likely reflects the models’ distinct parameterizations: the OU model’s regime-specific interactions (Surge × HL^+^ and Surge × HL^−^) may capture dynamics more specifically aligned with anxiety-related hyperarousal, whereas the React & Rebound model’s single interaction term captures a broader autonomic inflexibility pattern associated with elevated scores across multiple symptom measures.

### 3.9 Physical Activity Confound

Adding subject-mean accelerometer magnitude as a covariate to the React & Rebound model changed *R*^2^ by less than 0.002 and left all interaction coefficients essentially unchanged (Δ*β <* 0.03), indicating that physical activity does not explain the reported effects.

## 4 Discussion

### 4.1 Principal Findings

This work pursued a systematic comparison of computational approaches to extracting emotion regulation parameters from wearable HRV data. Several findings emerged. First, static metrics are insufficient: sympathetic load explains only 16% of variance, confirming that cumulative stress exposure alone does not capture vulnerability. Second, our attempt to measure discrete event restoration failed—decay rates from specific stress episodes showed no correlation with GAD-7 scores—challenging intuitions from laboratory paradigms. Third, both successful models require interaction terms; main effects alone are inadequate. The React & Rebound model with just three parameters (reactivity, rebound, and their interaction) achieves *R*^2^ = 0.60—nearly equivalent to a full model with sympathetic load and three-way interactions (*R*^2^ = 0.607)—while producing clearly significant coefficient estimates. The OU model achieves *R*^2^ = 0.69 with more sophisticated estimation. Importantly, these *R*^2^ values quantify the strength of association between autonomic dynamics and anxiety symptoms within the joint model, not out-of-sample prediction accuracy.

### 4.2 Regime Dynamics vs. Discrete Restoration

The failure of discrete event analysis is theoretically informative. Real-world emotional challenges are continuous and overlapping rather than discrete—what matters is not restoration after identifiable stressors but the probability of remaining in or exiting prolonged activation regimes. This finding is consistent with the perseverative cognition hypothesis (Brosschot et al., 2006), which posits that prolonged physiological activation persists not because of the stressor itself but because of sustained cognitive processing (worry, rumination) that maintains the stress response. Our regime-level framing— measuring transition probabilities rather than event-level restoration—may therefore be capturing the physiological signature of perseverative cognition, where it is the *system’s tendency to remain activated* that reflects maladaptive processing, not the return to calm after any specific event.

### 4.3 Autonomic, Circuit, and Cognitive Levels

The model parameters admit interpretation across levels: autonomically, reactivity and rebound describe sympathetic-parasympathetic dynamics; at the neural circuit level (Thayer and Lane, 2009), reactivity plausibly reflects amygdala responsivity while rebound reflects prefrontal regulatory capacity; cognitively, the interaction aligns with models emphasizing that anxiety symptoms emerge from threat sensitivity combined with regulatory failure—not either alone.

### 4.4 Implications for Digital Mental Health

The React & Rebound model has several properties that make it attractive for applied use. First, it requires only a consumer-grade wearable with PPG-based heart rate monitoring— devices already worn by millions of people. Second, the two-parameter framework (reactivity and rebound) is intuitive for both researchers and end users: “How easily is your nervous system activated?” and “How quickly does it calm?” Third, the autonomic profile quadrants (Table 5) suggest personalized intervention targets matched to each individual’s specific pattern. High reactivity reflects atypical salience network function (Menon and Uddin, 2010), in which the anterior insula assigns increased emotional salience to routine events, lowering the threshold for autonomic activation, and may respond to top-down cognitive strategies such as cognitive reappraisal, stress inoculation, or graded exposure training that raise the activation threshold. Slow rebound is characterized by prolonged time in sympathetically activated states, a pattern consistent with reduced parasympathetic re-engagement that plausibly maps to prefrontal-vagal regulatory capacity. Individuals displaying both high reactivity and slow rebound—the quadrant with the highest symptom severity—show a distinct autonomic profile, while those with high reactivity but fast rebound show symptom levels comparable to low-reactivity groups, suggesting that the two parameters capture separable dimensions of autonomic function.

In a consumer wearable application, the model could be implemented as follows: (1) an optional baseline questionnaire (e.g., GAD-7) at onboarding provides an informative prior that sharpens initial parameter estimates; (2) continuous HRV data from a smartwatch is processed into the autonomic balance signal; (3) transition probabilities are estimated from accumulated data, refining as more data arrives; and (4) the resulting autonomic profile—how readily the system enters activated states and how quickly it returns to baseline—is tracked over time, providing a continuous characterization of autonomic dynamics beyond static, aggregate measures.

### 4.5 Comparison to Existing Wearable Stress Metrics

Current wearable stress metrics from major platforms (Oura’s “Stress” score, Fitbit’s “Stress Management Score,” Apple Watch’s “Heart Rate Variability”) primarily rely on aggregate HRV measures—typically overnight RMSSD or time-domain metrics averaged over fixed windows. These approaches have two fundamental limitations that our model addresses. First, they collapse temporal dynamics into point estimates, treating a day with many brief stress responses (high reactivity, fast rebound) identically to a day with fewer but prolonged activations (low reactivity, slow rebound). Our results show these profiles are psychologically distinct. Second, existing metrics do not model interactions between reactivity and restoration, missing the key finding that reactivity alone is not associated with anxiety symptoms—only its combination with slow rebound.

The static sympathetic load metric we tested (*R*^2^ = 0.16) is conceptually similar to these commercial approaches. The React & Rebound model’s substantially stronger association with anxiety symptoms (*R*^2^ = 0.60) demonstrates the value of moving from static to dynamic autonomic modeling—capturing not just *how much* activation occurs, but *how the system transitions between states*.

### 4.6 Markov Assumption and Measurement Considerations

The React & Rebound model assumes that transition probabilities are stationary within a subject—that is, the probability of entering or staying in the activated state does not change systematically over the recording period. While this is a simplification (transition probabilities likely fluctuate with circadian rhythms, daily context, and longer-term mood changes), the strong association between transition parameters and trait-level GAD-7 scores provides indirect support for reasonable stationarity over the 4-week recording period. Future work should explore time-varying transition models that could capture within-person dynamics.

Additionally, the estimates of *P*(enter) and *P*(stay) carry posterior uncertainty. In the joint model, this uncertainty is naturally propagated through the hierarchical structure. However, in prospective applications where parameters are estimated from HRV data alone (without a concurrent GAD-7 score), measurement error in the transition probabilities would attenuate observed correlations, suggesting that the true associations between transition parameters and anxiety symptoms may be stronger than what would be observed in a prediction-only setting. Incorporating a baseline questionnaire at onboarding could help calibrate initial parameter estimates until sufficient HRV data accumulates.

### 4.7 Limitations

Several limitations warrant consideration. The sample (*n* = 44–49) comprised healthy young adults from a single region (South Korea), with most scoring in the low range of anxiety symptom severity. With *N* = 44, the study had approximately 80% power to detect correlations of *r* ≥ 0.41 at *α* = 0.05; replication in larger, more diverse samples spanning moderate-to-severe GAD-7 levels is needed. The cross-sectional design establishes correlation, not causation—longitudinal studies are needed to determine whether changes in transition parameters precede changes in anxiety symptoms. The HIGH/LOW state threshold, while grounded in the 90th percentile of the balance signal distribution, involves some arbitrariness, and consumer-grade PPG has lower precision than ECG. Although we showed that controlling for accelerometer-derived physical activity does not change the key findings (Δ*R*^2^ < 0.002), other potential confounds—caffeine, medication use, and contextual factors—remain uncontrolled. The Markov assumption of stationary transition probabilities is a simplification that should be relaxed in future work.

As noted in the Validation Approach, the reported *R*^2^ values (0.69 for OU, 0.60 for React & Rebound) reflect association strength within the joint model, not prospective prediction accuracy from HRV alone. We therefore emphasize converging validation evidence—outcome specificity analysis, robustness to physical activity confounds, and autonomic profile separation—rather than *R*^2^ alone as the basis for claiming utility. Independent replication with held-out datasets and prospective studies are needed to establish the model’s utility in practice.

Future work should incorporate multiple mental health instruments, ecological momentary assessment, systematic confound control, and development of efficient sequential estimation approaches for real-time application on consumer devices.

### 4.8 Conclusion

This work demonstrates that joint hierarchical Bayesian models can extract interpretable, psychologically meaningful emotion regulation parameters from naturalistic wearable data. The key finding—that elevated anxiety symptom severity is associated with the *interaction* between high reactivity and slow rebound—bridges autonomic dynamics measurable via consumer devices to neural circuit function to cognitive models of emotion regulation. The React & Rebound framework offers a parsimonious, interpretable parameterization with associations across multiple self-report symptom measures. By characterizing individual autonomic profiles—how readily the nervous system enters activated states and how quickly it returns to baseline—the framework provides a foundation for tracking these dynamics over time via consumer wearables.

## Data Availability

The wearable HRV dataset used in this study is publicly available from Baigutanova et al. (2025).

## Author Contributions

EE, CJ, and NG contributed to study conceptualization and manuscript review. ACH and TJS designed the computational framework, developed the models, conducted the analyses, and wrote the manuscript. AG provided scientific oversight and edited the manuscript.

## Multimedia Appendices

- **Multimedia Appendix 1:** MCMC trace plots for both models showing chain mixing and marginal posterior distributions.
- **Multimedia Appendix 2:** Posterior predictive checks comparing observed GAD-7 distribution to model predictions.
- **Multimedia Appendix 3:** PSIS-LOO diagnostics: Pareto *k* values per subject for both models.
- **Multimedia Appendix 4:** Full outcome specificity results (GAD-7, PHQ-9, ISI) across both models.
- **Multimedia Appendix 5:** Complete MCMC convergence diagnostics for all model parameters.

## Abbreviations

bpm: beats per minute
ESS: effective sample size
GAD-7: Generalized Anxiety Disorder 7-item scale
HDI: highest density interval
HR: heart rate
HRV: heart rate variability
IBI: inter-beat interval
ISI: Insomnia Severity Index
MCMC: Markov chain Monte Carlo
NUTS: No-U-Turn Sampler
OU: Ornstein-Uhlenbeck
PHQ-9: Patient Health Questionnaire 9-item scale
PPG: photoplethysmography
PSIS-LOO: Pareto-smoothed importance sampling leave-one-out
RMSSD: root mean square of successive differences
SEM: standard error of measurement
vmPFC: ventromedial prefrontal cortex

