## Supplementary Materials for "The React & Rebound Model: Capturing Emotion Regulation Dynamics from Passive Wearable Data"

#### Contents

1 Multimedia Appendix 1: MCMC Trace Plots 2

2 Multimedia Appendix 2: Posterior Predictive Checks 5

3 Multimedia Appendix 3: PSIS-LOO Diagnostics 7

4 Multimedia Appendix 4: Outcome Specificity 8

5 Multimedia Appendix 5: MCMC Convergence Diagnostics 10

#### Multimedia Appendix 1: MCMC Trace Plots

Trace plots showing chain mixing and marginal posterior distributions for both models. All four chains are overlaid; well-mixed chains appear as overlapping, stationary traces.

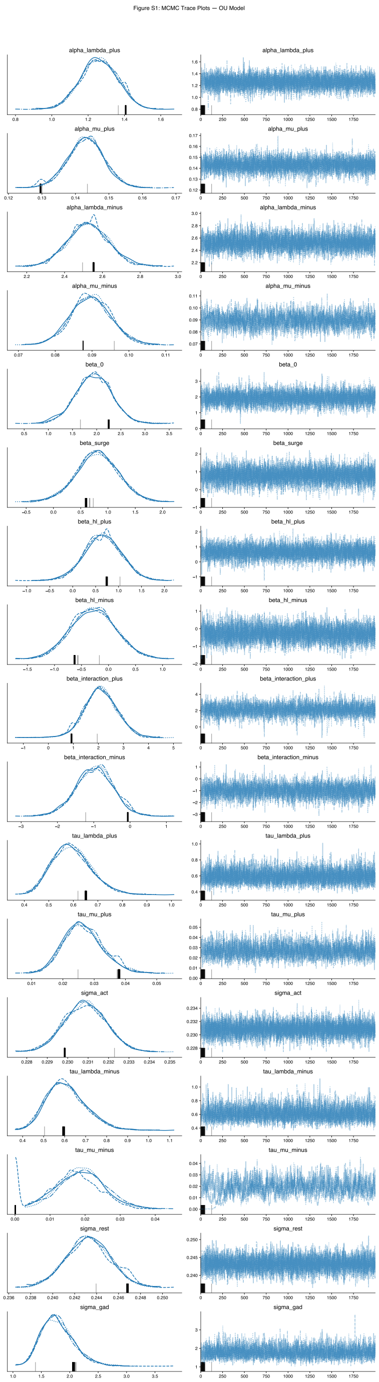


**Figure 1: OU model trace plots.** MCMC trace plots for the Ornstein–Uhlenbeck piecewise model regression coefficients and key hyperparameters. Left panels show posterior density estimates; right panels show trace plots across 4 chains × 2,000 post-warmup draws. All chains show good mixing with no visible trends or multimodality. Five divergent transitions were observed across 8,000 post-warmup draws (0.06%), well below the 1% threshold typically considered concerning.

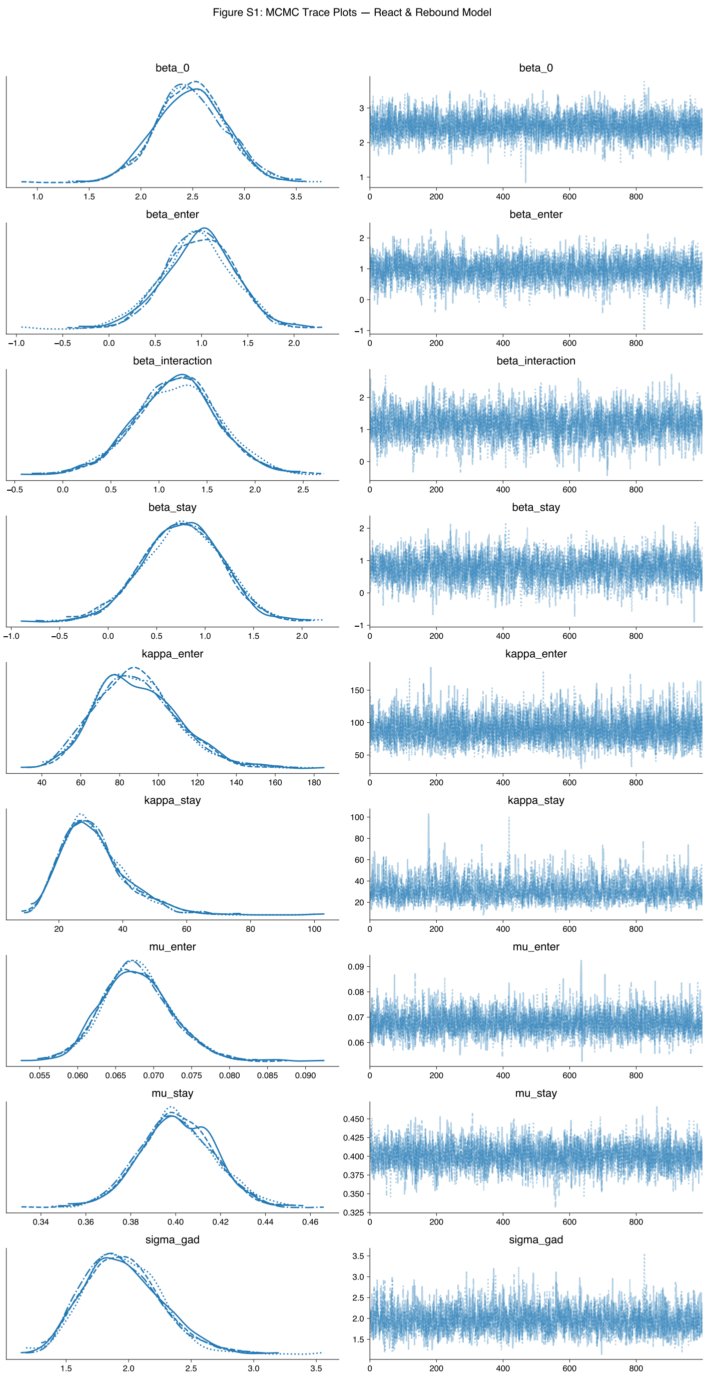


**Figure 2: React & Rebound model trace plots.** MCMC trace plots for the React & Rebound model regression coefficients and hierarchical hyperparameters ($\mu_{\text{enter}}$, $\kappa_{\text{enter}}$, $\mu_{\text{stay}}$, $\kappa_{\text{stay}}$). All chains show excellent mixing and convergence.

#### Multimedia Appendix 2: Posterior Predictive Checks

Posterior predictive checks compare the observed distribution of GAD-7 scores to distributions generated from the fitted model. Close agreement between observed and predicted distributions indicates adequate model fit.

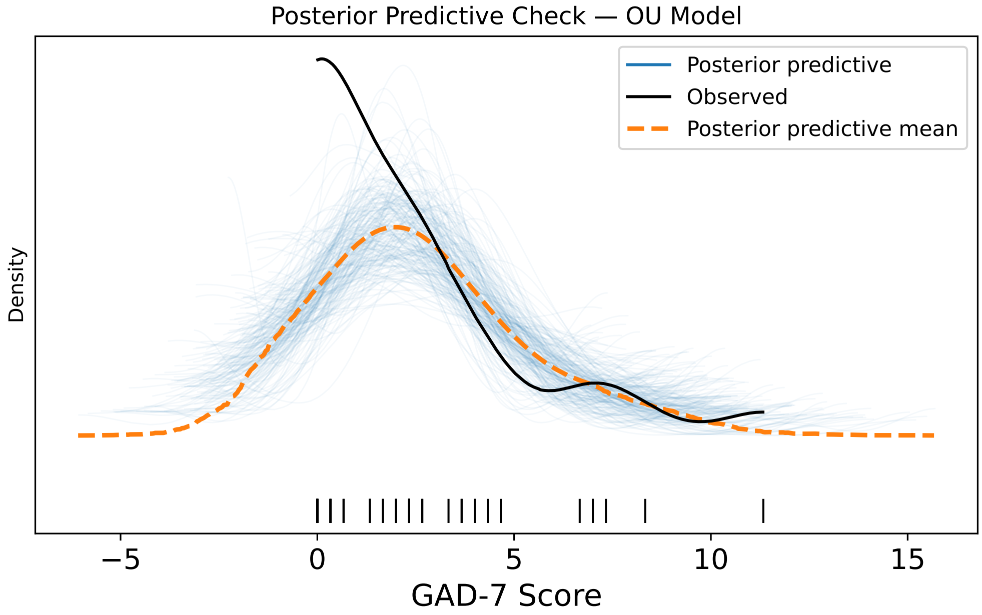


**Figure 3: OU model posterior predictive check.** The observed GAD-7 distribution (dark line) compared to 200 posterior predictive draws (light lines) and the posterior predictive mean (dashed line). The model captures the shape and spread of the observed distribution, including the right-skewed nature of GAD-7 scores in a predominantly healthy sample. Posterior predictive draws extend slightly below zero because both models use a Gaussian likelihood; a truncated Normal alternative was evaluated but produced inflated variance and wider credible intervals without improving coefficient estimation, so the standard Normal was retained.

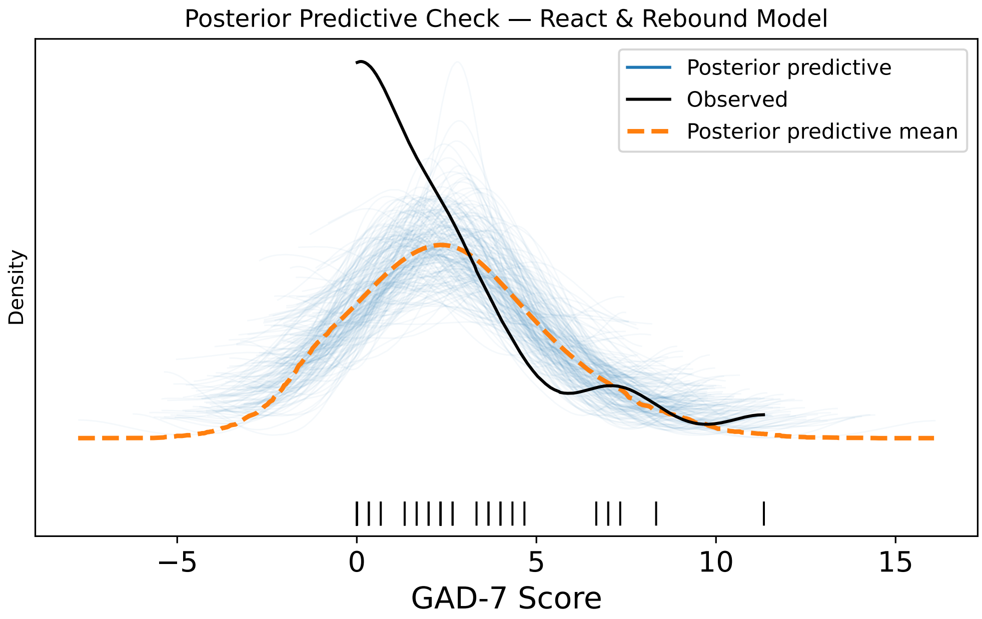


**Figure 4: React & Rebound model posterior predictive check.** The observed GAD-7 distribution (dark line) compared to 200 posterior predictive draws (light lines) and the posterior predictive mean (dashed line). Model fit is comparable to the OU model, adequately reproducing the observed distribution despite using fewer parameters.

### Multimedia Appendix 3: PSIS-LOO Diagnostics

Pareto-smoothed importance sampling leave-one-out (PSIS-LOO) cross-validation diagnostics identify observations that may exert disproportionate influence on the posterior. Pareto $k > 0.7$ flags potentially influential observations that warrant examination.

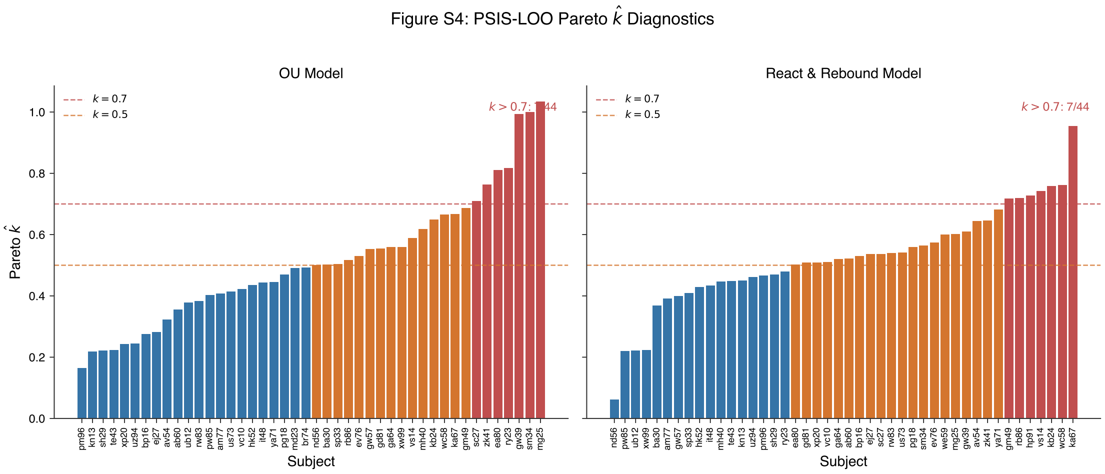


**Figure 5: PSIS-LOO diagnostics.** Pareto $k$ values for each subject in the OU model (left) and React & Rebound model (right). Dashed lines mark the $k = 0.7$ threshold. Seven subjects exceeded this threshold in each model (∼16% of the sample), consistent with expectations for datasets of this size ($N = 44$–$46$). Removing flagged subjects did not substantively change key interaction effects.

**Table 1:** PSIS-LOO summary statistics for both models.

| Model | ELPD-LOO | SE | Max Pareto $k$ | Subjects $k > 0.7$ |
|---|---|---|---|---|
| OU (Full) | −98.5 | 5.8 | 1.03 | 7/44 (16%) |
| React & Rebound (Full) | −100.3 | 5.5 | 0.95 | 7/45 (16%) |

#### Multimedia Appendix 4: Outcome Specificity

Full results of both models applied to all three self-report symptom measures: GAD-7 (anxiety symptoms), PHQ-9 (depression symptoms), and ISI (insomnia symptoms). For each model–outcome pair, a separate joint hierarchical Bayesian model was fit, and the posterior mean coefficients ($\beta$), 95% highest density intervals (HDI), and posterior probabilities are reported. $R^2$ values are Bayesian $R^2$, computed as 1 minus the ratio of residual to total sum of squares using posterior predictive means, consistent with the main text.

Permutation $p$-values provide a complementary frequentist significance test. For each outcome, we regressed the outcome scores on z-scored empirical Bayes estimates of the transition parameters (logit $P(\text{enter})$, logit $P(\text{stay})$, and their product) using OLS, and recorded the observed interaction coefficient. We then permuted the outcome vector 1,000 times, refitting the OLS regression each time, to generate a null distribution of interaction coefficients. The permutation $p$-value is the proportion of permuted $|\beta_{\text{int}}|$ values equal to or exceeding the observed $|\beta_{\text{int}}|$ (two-sided).

**Table 2:** Outcome specificity of the Ornstein–Uhlenbeck piecewise model.

| Outcome | Predictor | $\beta$ | 95% HDI | $P(\beta > 0)$ |
|---|---|---|---|---|
| **GAD-7 (anxiety symptoms)** | Surge | 0.81 | [0.05, 1.60] | 0.98 |
| | $HL^+$ | 0.66 | [−0.11, 1.48] | 0.95 |
| | $HL^-$ | −0.28 | [−1.11, 0.53] | 0.25 |
| | Surge × $HL^+$ | 2.08 | [0.87, 3.46] | 1.00 |
| | Surge × $HL^-$ | −0.97 | [−2.02, 0.00] | 0.03 |
| | $R^2 = 0.70$ | | Perm. $p = 0.009$ | |
| **PHQ-9 (depression symptoms)** | Surge | 0.77 | [−0.31, 1.83] | 0.92 |
| | $HL^+$ | 1.05 | [−0.10, 2.19] | 0.96 |
| | $HL^-$ | −0.22 | [−1.35, 0.98] | 0.36 |
| | Surge × $HL^+$ | 1.80 | [−0.45, 3.83] | 0.95 |
| | Surge × $HL^-$ | −1.12 | [−2.73, 0.45] | 0.08 |
| | $R^2 = 0.52$ | | Perm. $p = 0.183$ | |
| **ISI (insomnia symptoms)** | Surge | 0.95 | [−0.40, 2.20] | 0.93 |
| | $HL^+$ | 0.76 | [−0.86, 2.26] | 0.83 |
| | $HL^-$ | 0.26 | [−1.40, 1.74] | 0.63 |
| | Surge × $HL^+$ | 0.98 | [−1.21, 3.23] | 0.81 |
| | Surge × $HL^-$ | −0.27 | [−1.93, 1.25] | 0.37 |
| | $R^2 = 0.31$ | | Perm. $p = 0.310$ | |

In the OU model, the Surge $\times$ $HL^+$ interaction is strongly credible only for GAD-7 ($P(\beta > 0) = 1.00$, permutation $p = 0.009$). For PHQ-9 and ISI, the interaction shows weaker and non-significant effects.

**Table 3:** Outcome specificity of the React & Rebound model. Same format as above.

| Outcome | Predictor | $\beta$ | 95% HDI | $P(\beta > 0)$ |
|---|---|---|---|---|
| **GAD-7 (anxiety symptoms)** | Reactivity | 0.94 | [0.22, 1.69] | 0.99 |
| | $\text{Rebound}^{-1}$ | 0.66 | [−0.15, 1.46] | 0.95 |
| | React. $\times$ $\text{Reb.}^{-1}$ | 1.26 | [0.35, 2.31] | 1.00 |
| | $R^2 = 0.61$ | | Perm. $p = 0.033$ | |
| **PHQ-9 (depression symptoms)** | Reactivity | 0.92 | [−0.13, 1.89] | 0.96 |
| | $\text{Rebound}^{-1}$ | 0.97 | [−0.14, 2.05] | 0.96 |
| | React. $\times$ $\text{Reb.}^{-1}$ | 1.52 | [0.14, 2.89] | 0.98 |
| | $R^2 = 0.53$ | | Perm. $p = 0.117$ | |
| **ISI (insomnia symptoms)** | Reactivity | 1.36 | [0.20, 2.46] | 0.99 |
| | $\text{Rebound}^{-1}$ | 1.27 | [0.09, 2.55] | 0.98 |
| | React. $\times$ $\text{Reb.}^{-1}$ | 1.97 | [0.35, 3.57] | 0.99 |
| | $R^2 = 0.58$ | | Perm. $p = 0.024$ | |

The two models show different specificity patterns. In the OU model, the activation–restoration interaction is strongly specific to GAD-7, with the PHQ-9 and ISI interactions falling short of conventional credibility thresholds. In the React & Rebound model, the reactivity $\times$ rebound interaction is credible across all three symptom measures ($P(\beta > 0) \geq 0.98$), though the OLS permutation test remains non-significant for PHQ-9 ($p = 0.117$). This difference likely reflects the models' distinct parameterizations: the OU model's regime-specific interactions (Surge $\times$ $HL^+$ and Surge $\times$ $HL^-$) may capture dynamics more specifically aligned with anxiety-related hyperarousal, whereas the R&R model's single interaction term captures a broader autonomic inflexibility pattern associated with elevated scores across multiple symptom measures.

#### Multimedia Appendix 5: MCMC Convergence Diagnostics

Complete convergence diagnostics for key model parameters. All parameters achieved $\hat{R} < 1.01$ with adequate effective sample sizes and zero divergent transitions.

**Table 4:** Posterior summaries and convergence diagnostics for regression coefficients and hyperparameters. $ESS_{bulk}$ = bulk effective sample size; $\hat{R}$ = split-$\hat{R}$ convergence statistic (values < 1.01 indicate convergence). Both models were estimated with 4 chains × 2,000 post-warmup draws (1,000 tuning draws discarded), target acceptance rate 0.95.

| Model | Parameter | Mean | SD | 95% HDI | $ESS_{bulk}$ | $\hat{R}$ |
|---|---|---|---|---|---|---|
| OU model | $\beta_0$ (Intercept) | 1.94 | 0.37 | [1.25, 2.63] | 6886 | 1.001 |
| | $\beta_{surge}$ (Surge frequency) | 0.81 | 0.39 | [0.05, 1.52] | 5822 | 1.000 |
| | $\beta_{HL^+}$ (Half-life, activated) | 0.66 | 0.41 | [−0.13, 1.42] | 4849 | 1.000 |
| | $\beta_{HL^-}$ (Half-life, rest) | −0.26 | 0.40 | [−0.99, 0.52] | 5577 | 1.000 |
| | $\beta_{int^+}$ (Surge × $HL^+$) | 2.09 | 0.66 | [0.85, 3.36] | 2536 | 1.001 |
| | $\beta_{int^-}$ (Surge × $HL^-$) | −1.00 | 0.52 | [−1.99, −0.03] | 2644 | 1.000 |
| | $\sigma_{GAD}$ | 1.78 | 0.28 | [1.29, 2.33] | 3261 | 1.000 |
| React & Rebound | $\beta_0$ (Intercept) | 2.35 | 0.35 | [1.71, 3.04] | 6452 | 1.000 |
| | $\beta_{enter}$ (Reactivity) | 0.92 | 0.38 | [0.22, 1.63] | 5057 | 1.000 |
| | $\beta_{stay}$ (Rebound, inverse) | 0.69 | 0.43 | [−0.11, 1.50] | 3618 | 1.001 |
| | $\beta_{int}$ (React. × Rebound) | 1.29 | 0.50 | [0.37, 2.23] | 2855 | 1.001 |
| | $\mu_{enter}$ | 0.072 | 0.005 | [0.063, 0.080] | 6497 | 1.001 |
| | $\kappa_{enter}$ | 90.3 | 19.5 | [55.7, 128.5] | 5758 | 1.000 |
| | $\mu_{stay}$ | 0.438 | 0.014 | [0.411, 0.464] | 5057 | 1.000 |
| | $\kappa_{stay}$ | 50.0 | 14.9 | [25.1, 78.2] | 2843 | 1.000 |
| | $\sigma_{GAD}$ | 1.99 | 0.31 | [1.44, 2.59] | 3040 | 1.000 |

For both models, all $\hat{R}$ values were below 1.01 and effective sample sizes (ESS) exceeded 400 for all regression parameters. No divergent transitions were observed. Subject-level parameters (44 $\lambda^+$, $\lambda^-$, $\mu^+$, $\mu^-$ parameters for the OU model; 46 $P(\text{enter})$ and $P(\text{stay})$ parameters for the React & Rebound model) all converged similarly. Trace plots for key parameters are shown in Multimedia Appendix 1.
